# TMPRSS2 and ADAM17 interactions with ACE2 complexed with SARS-CoV-2 and B^0^AT1 putatively in intestine, cardiomyocytes, and kidney

**DOI:** 10.1101/2020.10.31.363473

**Authors:** Bruce R. Stevens

## Abstract

COVID-19 outcomes reflect organ-specific interplay of SARS-CoV-2 and its receptor, ACE2, with TMPRSS2 and ADAM17. Confirmed active tropism of SARS-CoV-2 in epithelial cells of intestine and kidney proximal tubule, and in aging cardiomyocytes, capriciously manifests extra-pulmonary organ-related clinical symptoms in about half of COVID-19 patients, occurring by poorly understood mechanisms. We approached this knowledge gap by recognizing a clue that these three particular cell types share a common denominator kindred of uniquely expressing the SLC6A19 neutral amino acid transporter B^0^AT1 protein (alternatively called NBB, B, B^0^) serving glutamine and tryptophan uptake. B^0^AT1 is a cellular trafficking chaperone partner of ACE2, shown by cryo-EM to form a thermodynamically-favored stabilized 2ACE2:2B^0^AT1 dimer-of-heterodimers. The gut is the body’s site of greatest magnitude expression depot of both ACE2 and B^0^AT1. This starkly contrasts with pulmonary pneumocyte expression of monomeric ACE2 with conspicuously undetectable B^0^AT1. We hypothesized that B^0^AT1 steers the organ-related interplay amongst ACE2, TMPRSS2, ADAM17, and SARS-CoV-2 RBD. The present study employed molecular docking modeling that indicated active site catalytic pocket residues of TMPRSS2 and ADAM17 each formed bonds ≤ 2 A with monomer ACE2 specific residues within a span R652-D713 involved in cleaving sACE2 soluble ectodomain release. These bonds are consistent with competitive binding interactions of experimental anti-SARS-CoV-2 drug small molecules including Camostat and Nafamostat. Without B^0^AT1, ACE2 residues K657 and N699 dominated docking bonding with TMPRSS2 or ADAM17 active sites, with ACE2 R710 and R709 contributing electrostatic attractions, but notably ACE2 S708 never closer than 16-44 A. However, in the dimer-of-heterodimers arrangement all ACE2 neck region residues were limited to TMPRSS2 or ADAM17 approaches 35 A, with the interference directly attributed to the presence of a neighboring B^0^AT1 subunit complexed to the partnering ACE2 subunit of 2ACE2:2B^0^AT1; ADAM17 failed to dock by bumping its active site pocket oriented dysfunctionally outwardly facing 180^0^ away. Results were the same whether the dimer-of-heterodimers was in either the “closed” or “open” conformation, or whether or not SARS-CoV-2 RBD was complexed to ACE2. The results implicate B^0^AT1-and in particular the 2ACE2:2B^0^AT1 complex-as a maJor player in the landscape of COVID-19 pathophysiology engaging TMPRSS2 and ADAM17, consistent with experimental evidence in the literature and in clinical reports. These findings provide a gateway to understanding the roles of B^0^AT1 relating to COVID-19 manifestations putatively assigned to intestinal and renal epithelial cells and cardiomyocytes, with underpinnings useful for considerations in public hygiene policy and drug development.

## INTRODUCTION

SARS-CoV-2 has been demonstrated to actively and productively infect human small intestinal dividing and post-mitotic enterocytes, infect cardiomyocytes, and appear as virion particles in kidney proximal tubule cells of COVID-19 patients [1-5]. These observations are consistent with findings that: i) nearly half of COVID-19 patients present with gastrointestinal (GI) symptomology as a risk factor [6-8]; ii) ∼6.5×10^2^ to 1.6×10^5^ active virion particles per day are reportedly shed in feces [6] with RNA detectable in toilet aerosols [7-11], even in SARS-CoV-2 positive subJects without pulmonary symptoms; iii) myocardial damage appears to correlate with outcome in autopsies showing myocarditis with SARS-CoV-2 viral RNA in hearts of COVID-19 patients; and iv) Hartnup-like neutral amino aciduria accompanies impaired renal proximal tubule function [1-3, 12]. In addition to these individual organ involvements, we [13-15] have posited a gut-lung axis integrative coupling of microbiome dysbiosis, disruption of tryptophan signaling, and systemic inflammasome events of COVID-19.

In spite of clinical prevalence and tissue tropisms, the mechanism is currently not known how some patients are spared yet others manifest acute or latent cardiovascular, GI, or renal comorbidity beyond known pulmonary pneumocyte involvement in COVID-19. Furthermore, hygiene policy concerning fecal to oral transmission, and COVID-19 drug development targeting GI, cardiac, and nephron involvement have been impeded by the lack of mechanistic insight.

A significant clue in this knowledge gap lies in the under-recognition that these three cell types in particular-enterocytes, renal proximal tubule epithelia, and aging (but not in young) cardiomyocytes-are nearly unique among all cell types of the body in expressing the components of a thermodynamically favored state of angiotensin converting enzyme 2 (ACE2) that is stabilized in a dimer-of-heterodimers complexed with B^0^AT1 neutral amino acid transporter [5, 13, 14, 16-22]. B^0^AT1 (alternately called NBB, B, B^0^) was previously originally discovered and functionally characterized by Stevens and coworkers [21, 23-29] as the sodium-coupled neutral amino acid transport system in intestinal epithelial cell apical brush border membranes[21], notably serving tryptophan and glutamine uptake. Subsequently, cloning of its SLC6A19 gene by Ganapathy, Broer, Fairweather, Verrey and colleagues was expanded to intimate epithelial cell chaperoning of B^0^AT1 by ACE2 [16, 18-20, 30-33].

The expression patterns ascertained by single cell RNA seq analyses, immunohistochemistry, and functional genomics studies demonstrate that the human GI tract is the body’s site of greatest magnitude expression of ACE2 and B^0^AT1-orders of magnitude greater than lung or any extra-GI tissues-with signficiant but lesser expressions in kidney [34-41]. Lung pneumocyte ACE2 appears to be expressed as the stand-alone monomer [13, 14, 42, 43]. The atomic structure assemblage of two ACE2 subunits plus two B^0^AT1subunits is organized as a 2ACE2:2B^0^AT1 dimer-of-heterodimers complex, as recently determined by Yan and coworkers in Zhou’s group [17] using cryo-electron microscopy, who reported hinging between “open” and “closed’ conformation states. However, the role of ACE2’s structure in steering COVID-19 events is not known, whether as a monomer, homodimer, or multimer complexed with B^0^AT1.

At the big picture level, it is well recognized that ACE2, transmembrane serine protease-2 (TMPRSS2), and sheddase disintegrin and metalloproteinase-17 (ADAM17) are enzymes that triangulate among themselves, and behave individually, as central players in SARS-CoV-2 infections and COVID-19 outcomes [13-15, 42-48]. All three enzymes are expressed in enterocytes, cardiomyocytes and proximal tubule [34-40]. Biochemical studies of these type I integral membrane proteases, in conJunction with ACE2 mutant/chimeric *in vitro* expression experiments, have established that ADAM17 and TMPRSS2 compete for ACE2’s involvement with SARS-CoV-2 through intertwined venues [13-15]: i) they tightly govern the potential for either limiting or promoting spiraling positive feedback dysregulation of the renin-angiotensin system (RAS) through ACE2’s beneficial pleiotropic peptidyl carboxypeptidase activity associated with cardiopulmonary, gastrointestinal and renal physiology; ii) they can either potentiate or limit the reactive oxygen species (ROS)/cytokine storm implicated in inflammation-mediated tissue damage and pathophysiology of COVID-19; iii) they steer the relative infectivity of SARS-CoV-2 by shedding soluble ACE2 ectodomain (sACE2) from the cell surface; and iv) control priming of SARS-CoV-2 S-protein for cell entry when ACE2 is physically anchored to the cell surface. The ACE2 ectodomain region is highJacked in COVID-19 as the receptor for the SARS-CoV-2 spike (S) protein receptor binding domain (RBD). TMPRSS2 not only primes the SARS-CoV-2 S-protein for cell entry, but putatively cleaves ACE2 to release sACE2 ectodomain into the extracellular milieu and away from protecting against the pernicious effects the other RAS components of cells. While it is known that ACE2 proteolysis by TMPRSS2 is not required per se for SARS-CoV-2 cell entry, it is clear that TMPRSS2 priming of the SARS-CoV-2 S protein is an obligatory step for virus cell entry and development of COVID-19 [13, 14, 42, 43, 46-48]. SARS-CoV-2 infectivity in human cells can be inhibited by the TMPRSS2 inhibitors Camostat, Nafamostat and several other experimental small molecules[49-52]. Interactions of TMPRSS2 with the parent ACE2 structure are not settled [53], other than a known integrin-binding motif involved with SARS-CoV-2 RBD attachment[54], and controversies exist whether anti-ADAM17 drugs targeting its HELGHNFGAEHD catalytic pocket motif would be protective or harmful[55].

In the light of the existing literature, we hypothesized that TMPRSS2 and ADAM17 engagement with ACE2 involves a consensus of residues within the of neck region ACE2 relating to a cleavage motif requisite for ectodomain shedding and/or S-protein processing, such that B^0^AT1 of the 2ACE2:2B^0^AT1 dimer-of-heterodimers complex structure in either “open” or “closed” conformations governs TMPRSS2 or ADAM17 access to ACE2, in contrast to ready access in monomeric ACE2, homodimeric ACE2:ACE2, or heterodimeric 1ACE2:1B^0^AT1 states. Our prior preliminary report [56] provided impetus for the current undertaking.

## METHODS

Protein-protein interaction software ClusPro 2.0 [57-59] was employed for molecular docking simulations which involved 70,000 rotations probing rigid body docking, clustering of lowest energy complexes, and final energy minimization. The docked complexes were scored based on energy considerations of the general form:

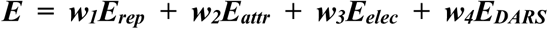

where *E*_*rep*_ is energy of repulsion from van der Waals interactions, *E*_*attr*_ is energy of attraction from van der Waals interactions, *E*_*elec*_ is the electrostatic energy term, and *E*_*DARS*_ is the energy of desolvation [59]. The *w* terms represent the weight of each coefficient depending on the scoring mode. Docking scoring was based on the “Balanced” option choice with constants *w*_*1*_ = 0.4, *w*_*2*_ = −0.4, *w*_*3*_ = 600, *and w_4_* = 1.0. After scoring, the top 1000 docked positions, based on score, were clustered together by finding a local center with the most docking neighbors within a 9A sphere [59]. This step was repeated to find multiple clustering centers. Finally, energy minimization removed any steric clashes between side chains, with the top docking scores outputted along with the corresponding complexed structure coordinate files [59]. Prior to docking, structures in the PDB files were modified by removing small molecules including solvent, inhibitors, ligands, glycosylation molecules, diacylglycerol 3-phosphate, and non- standard amino acids. For each docking simulation, non-physiologically relevant repulsion domains were masked out using PyMOL [60] to prevent non-specific dockings (e.g., unrealistic access of TMPRSS2 or ADAM17 ligand binding pockets to the hydrophobic residues embedded within the transmembrane domains of ACE2 and/or B^0^AT1 membrane anchors). For each paired docking of ligand chain with receptor chain, the Cluster 0 set of residues garnering the greatest clustering and most negative docking energy score was assessed for interface contact distances using ChimeraX software [61] meeting default probe criteria 1.4 A or being buried with a 15 A^2^ area cutoff; such interface contact distances were typically ≤ 2.0 A. Figures were generated using PyMOL [60] and ChimeraX [61].

PDB ID:6M17 was employed as the 2ACE2:2B^0^AT1/SARS-CoV-2 receptor binding domain (RBD) complex receptor in the “closed” conformation; “closed” conformation PDB ID:6M18 was employed analogously lacking SARS-CoV-2 RBD. PDB ID:6M1D was employed as the “open” conformation of 2ACE2:2B^0^AT1 dimer-of-heterodimers. Monomers, homodimers, heterodimers, and tetramers were derived by parsing chains of 6M17, 6M18, or 6M1D using PDBEditor [62] and PyMOL v2.4.0 [60]. Although 1ACE2:1B^0^AT1 heterodimers do not have physiological antecedence with experimental evidence, these dockings were included to complete the combination permutations of ACE2 and B^0^AT1 subunit interactions. The receptor attraction set employed was specified as ACE2 residues R652, Q653, Y654, F655, L656, K657, V658, K659, R708, S709 chosen based on TMPRSS2 and ADAM17 experimental cleavage studies [45, 47, 48, 53]. For dimer-of-heterodimers and homodimer dockings, modeling was conducted with these residues either singly exposed on one ACE2 chain and masked on the other chain, or dually exposed on both ACE2 chains.

The TMPRSS2 ligand employed was the well-studied SWISS-MODEL [63] Repository (UniProtKB accession O15393) human isoform-2 [49-51, 64] built on the PBD ID: 5CE1.1.A template of human serine protease hepsin complex, with crystallography inhibitor removed. This was used because the complete atomic structure of TMPRSS2 is not available in the literature. We selected TMPRSS2 ligand attraction residues as H296, D345, D435, S436, C437, Q438, S441, T459, S460, W461, C465, V473, and Y474 which incorporates the catalytic and substrate binding motifs (residue numbering convention of Swiss-Model 015393). Ramachandran plot validation of the overall structure and placement of active site residues was conducted using MolProbity ver. 4.4 software [65] in conJunction with SWISS-MODEL workspace [63].

ADAM17 ligand was chosen as the monomer chain A of PDB ID:3LGP, with the small molecule inhibitor removed. The docking attractor residues were chosen as H405, E406, L407, G408, H409, N410, F411, G412, A413, E414, H415, and D416 which incorporated zinc atom coordinating active site residues of H405, H409, H415 and E406 [66].

## RESULTS

The TMPRSS2 ligand modeled on PDB ID: 5CE1.1.A template was validated [63, 65] as shown in the Ramachandran plot of Fig. 1. There were 94.24% favored rotamers, notably including favorable placement of the known active site pocket catalytic triad H296, ASP345 and S441 (red circles), with 0.87% outliers R225, A216 and S208 (black circles) far from the active site residues, which are not reported to be involved either catalytically nor with meaningful impact on conformation [63, 65]. Overall structure QMEAN = −1.42.

**Fig. 1.**
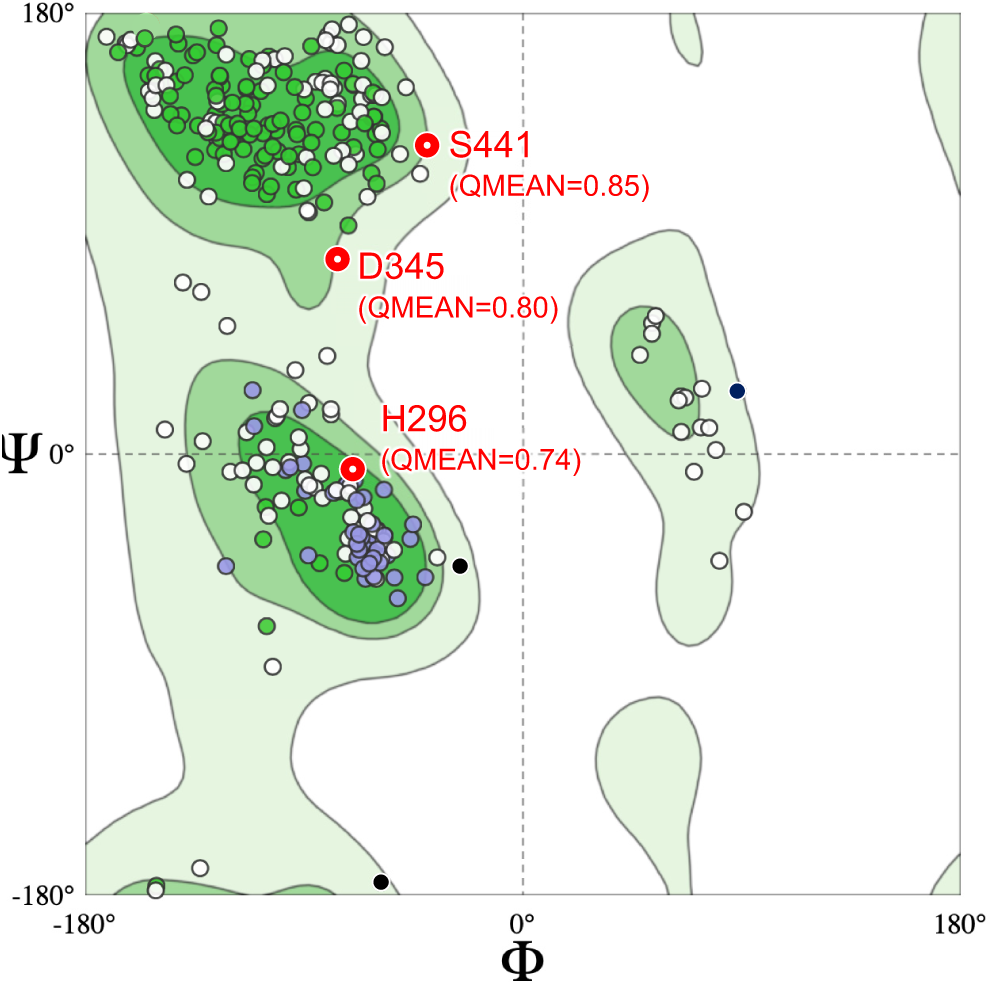
Ramachandran plot validating TMPRSS2 ligand structure employed in the present study. TMPRSS2 ligand structure modeled on PDB ID: 5CE1.1.A template showing favorable placement of active site pocket catalytic triad (red circles)

The atomic structure of the ACE2:B^0^AT1 dimer-of-heterodimers has been reported for two conformation states-”closed” (PBD ID:6M18 without SARS-CoV-2 RBD, and 6M17 with SARS-CoV-2),and “open” (PDB ID:6M1D) [17]. While the ACE2 ectodomain head is reported to move between these states [17], the movement distances within the hinged neck region itself have not been reported. Fig. 2A shows the overall organization of the dimer-of-heterodimers anchored in the plasma membrane in these two states emphasizing the hinging neck region, with the closeup in Fig. 2B and 2C detailing movements of the particular motif of residues of interest in the present study. The overall median excursion distance limited to shifting within the neck region is roughly 11 A (movement 5.5 A left plus 5.5 A right), as experienced by docking TMPRSS2 or ADAM17 ligands approaching from the front (2B) or top (2C) perspectives.

**Fig. 2.**
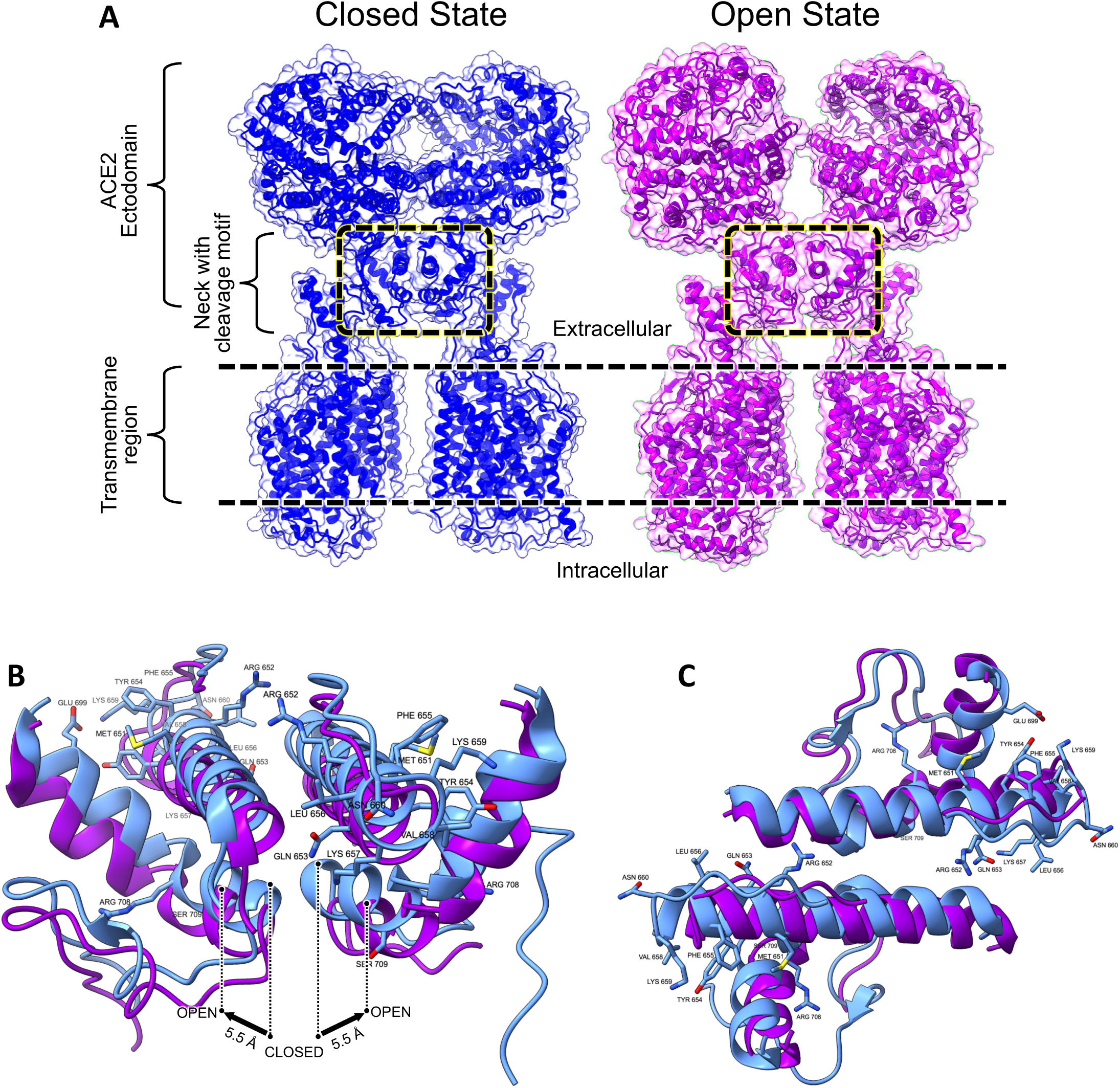
Closed vs. Open states of 2ACE2:2B0AT1 dimer-of-heterodimers complex encountered by TMPRSS2 or ADAM17. (A) Two ACE2 subunits plus two B^0^AT1 subunits complexed as a dimer of two ACE:B^0^AT1 heterodimers in the “CLOSED” conformation state (PBD ID:6M18, blue) and “OPEN” state (PBD ID:6M1D, purple), as observed by cryo-electron microscopy [17]. The complex undergoes movement between these states by pivoting at the neck region. Both neck region boxes from (A) are superpositioned and enlarged in the front view (B) and top view (C). The complex is anchored within the plasma membrane. (B) Docking target residue motif in neck region in the closed (blue) or open (purple) states, showing front approach perspective encountered as seen by TMPRSS2 or ADAM17 active site pockets. Overall median excursion distance shifted within the neck region is roughly 11 A (movements 5.5 A left plus 5.5 A right). (**C)** Docking target residue motif looking from the top perspective onto docking target motif of ACE2 residues showing “closed” (blue) and “open” (purple) states.

Fig. 3 shows successful docking of TMPRSS2 to stand-alone ACE2 monomer complexed with SARS-CoV-2 RBD, with details reported in Table 1. Residues of TMPRSS2 and ACE2 meeting criteria for interfacing “contact” are shown in Fig. 3B, demonstrating participation of the ACE2 neck region motif residues of Fig. 2B and 2C. An exploded separation of this contact zone emphasizing the active site catalytic pocket residues of TMPRSS2 is shown in the closeup in Fig. 3C, whereby select contact bond distances are indicated within esthetic graphical limitations, with all contact bond distances reported in Table 1.

**Table 1.**
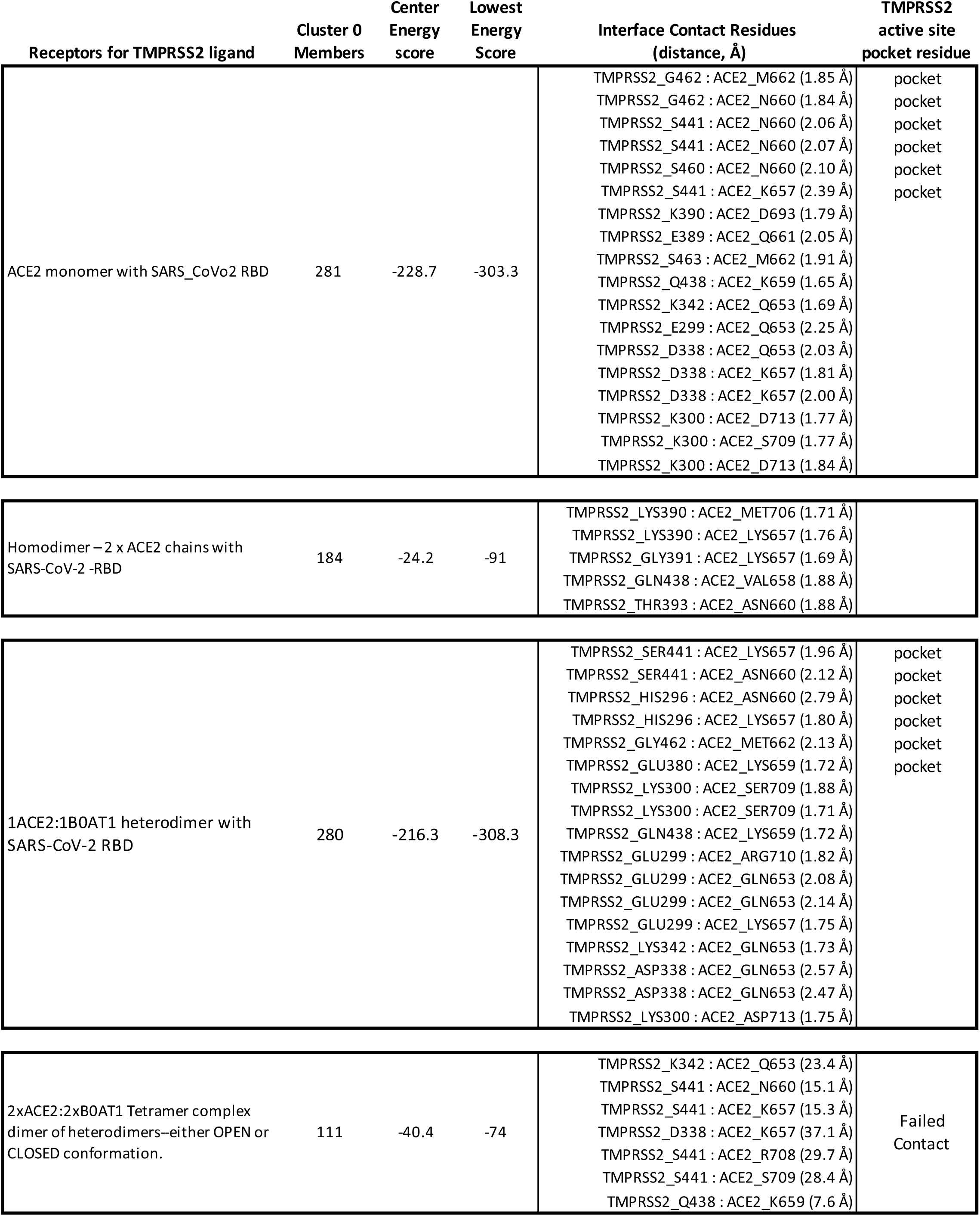
TMPRSS2 ligand molecular docking contact residues with various ACE2 structures.

**Fig. 3.**
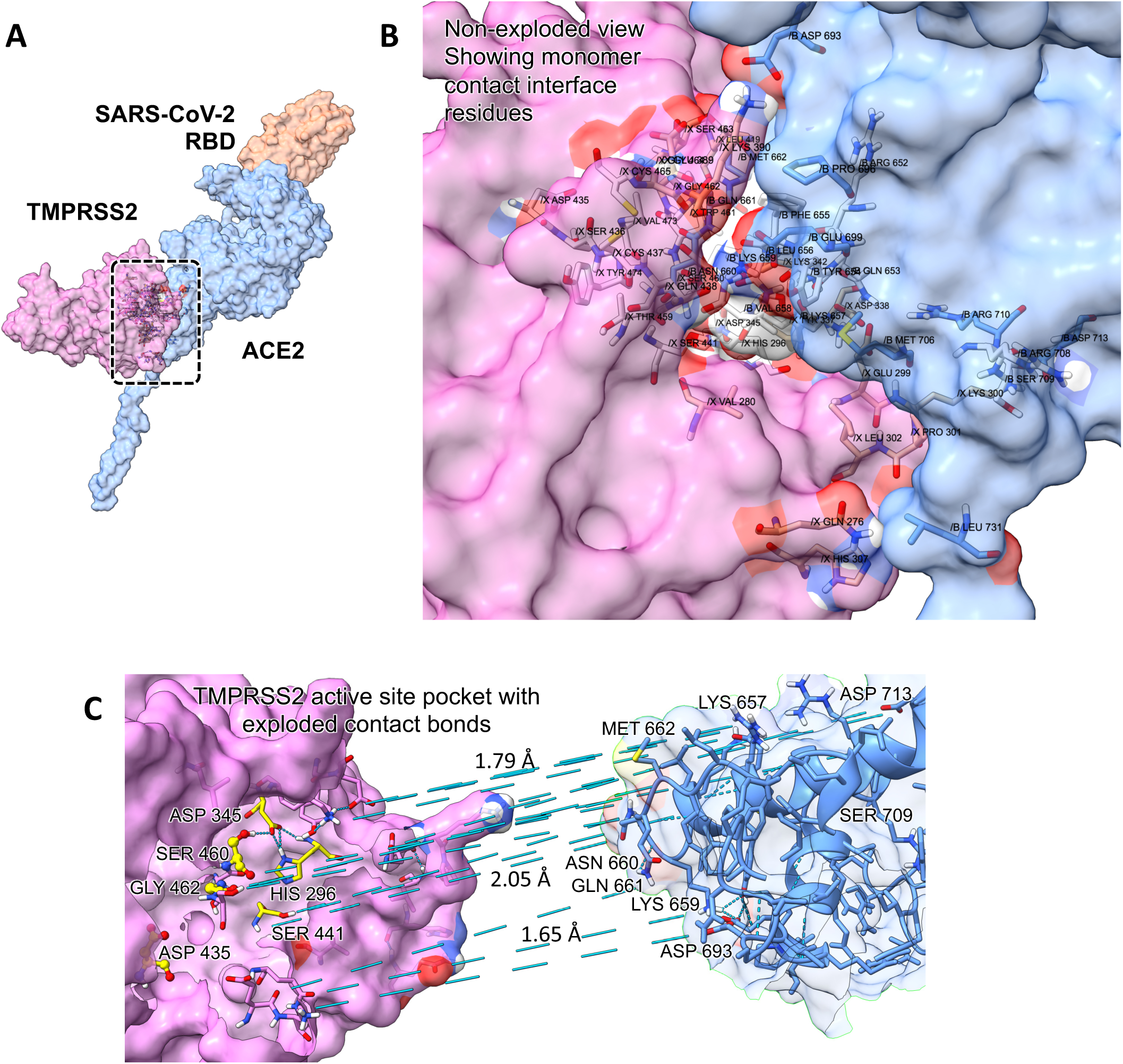
TMPRSS2 docked with ACE2 monomer complexed with SARS-CoV-2 RBD. (A) Docking model results showing ACE2 monomer (light blue), TMPRSS2 (pink), and SARS-CoV-2 RBD (gold). (B) Closeup of box in (A) showing successful docking interface contact residues of TMPRSS2 catalytic active site pocket closely engaged with ACE2 ectodomain putative cleavage zone residues. (C) Exploded native view of (B) showing TMPERSS2 active site pocket and select interface contact distances to ACE2 as measured using pre-exploded distances. Strategic pocket components are catalytic residues (yellow) H296, K345 and S441, and substrate binding residues D435, S460 and G462. Interaction details are in Table 1.

TMPRSS2 docked with ACE2:ACE2 homodimer whether complexed with SARS-CoV-2 RBD (Fig. 4) or not including RBD (not shown). The same model results summarized in Table 1 were obtained whether docking conditions were restricted to availability of one or simultaneously both ACE2 subunit attractor residues. An exploded view of the docking interface contact residues is shown in Fig. 4B along with select distances, with Table 1 reporting all bonds and distances. Although docking involved contact with five TMPRSS2 residues, none of those bonds involved an active site pocket residue.

**Fig. 4.**
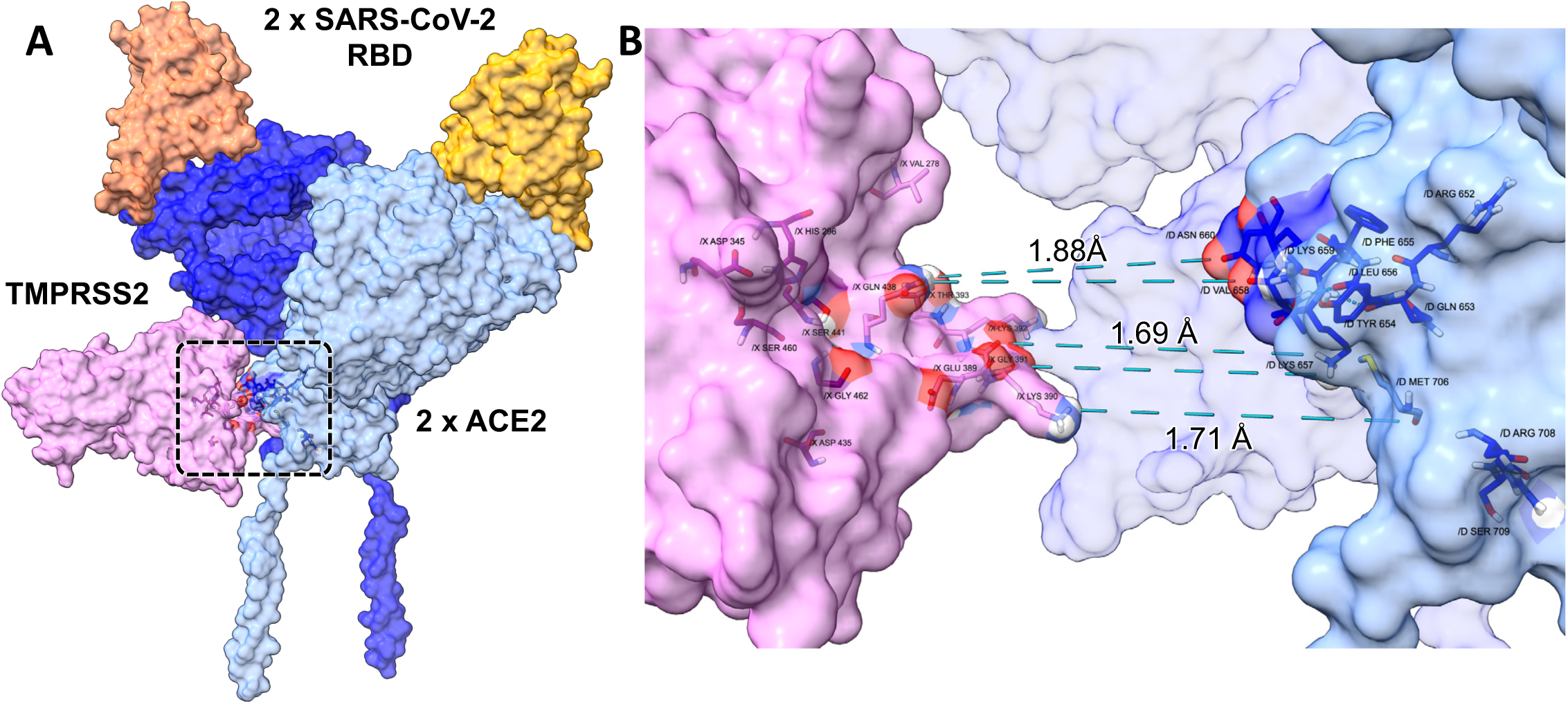
TMPRSS2 docked with ACE2:ACE2 homodimer complexed with SARS-CoV-2 RBD. (A) Docking model results showing ACE2 homodimer (light and dark blue), TMPRSS2 (pink), and SARS-CoV-2 RBD (dark and light gold). The same model results were obtained whether docking conditions were restricted to availability of one or simultaneously both ACE2 subunit attractor residues. (B) Exploded view of box in (A) showing docking interface contact residues of TMPRSS2 catalytic active site pocket engaged with ACE2 ectodomain cleavage zone residues. Select interface contact distances are shown, as measured using pre-exploded distances. The same model results were obtained whether docking conditions were restricted to availability of one or simultaneously both ACE2 subunit attractor residues. Interaction details are in Table 1.

The active site pocket of TMPRSS2 docked successfully with a heterodimer comprised of 1ACE2:1B^0^AT1 complexed with SARS-CoV-2 RBD, as shown in Fig. 5. An exploded view of the docking interface contact residues is shown in Fig. 5B along with select distances, with Table 1 reporting all bonds and distances.

**Fig. 5.**
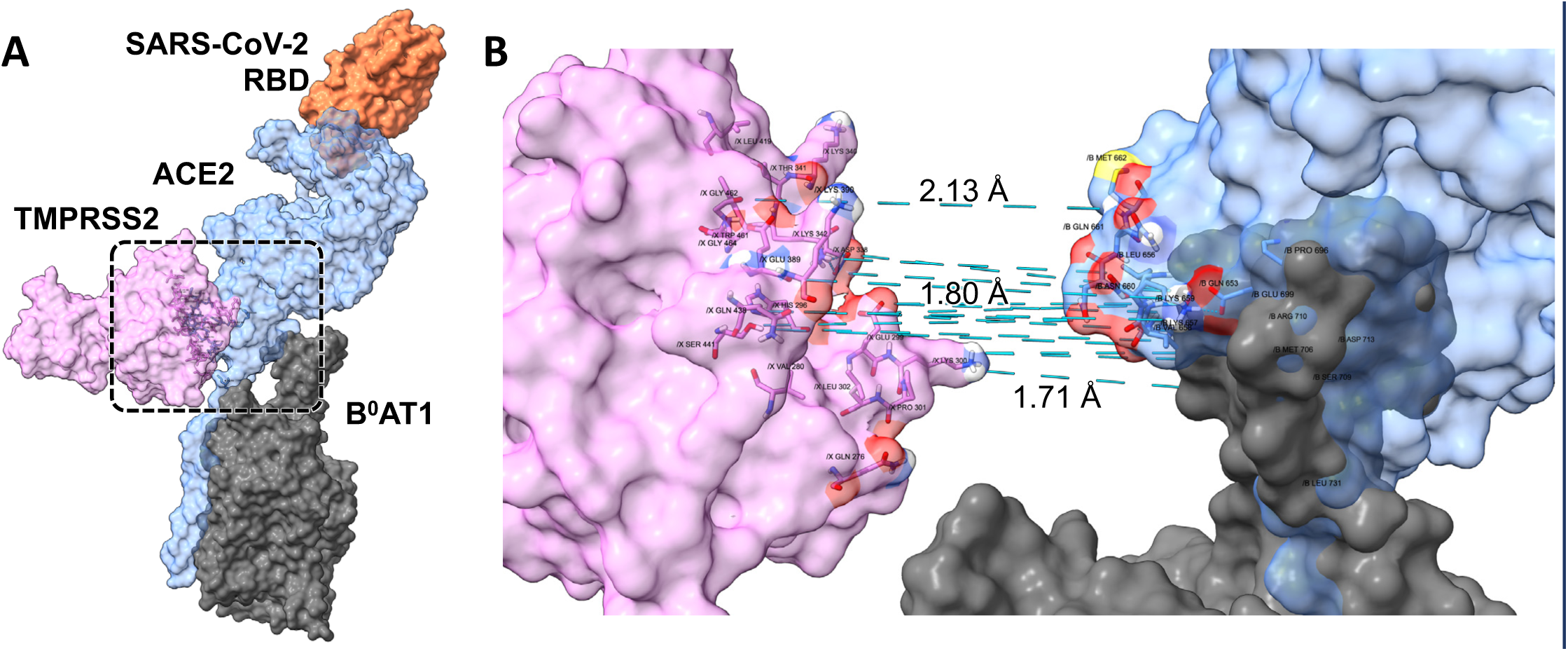
TMPRSS2 docked with 1ACE2:1BOAT1 heterodimer complexed with SARS-CoV-2 RBD. (A) Docking model results showing heterodimer ACE2 (light blue), B^0^AT1 (gray), TMPRSS2 (pink), and SARS-CoV-2 RBD (gold). (B) Exploded view of box in (A) showing docking interface contact residues of TMPRSS2 catalytic active site pocket closely engaged with ACE2 ectodomain cleavage zone residues. Select interface contact distances are shown, as measured using pre-exploded distances. The same model results were obtained whether docking conditions were restricted to availability of one or simultaneously both ACE2 subunit attractor residues. Interaction details are in Table 1.

Figure 6 and the data of Table 1 indicated that TMPRSS2 ligand failed to dock with the closed state 2ACE2:2B^0^AT1 dimer-of-heterodimers neck region, whether or not the ACE2 was complexed to SARS-CoV-2 RBD. Instead, TMPRSS2 randomly stuck to the dimer-of-heterodimers in orientations and distances generally ∼15-30 A from TMPRSS2 active site pocket residues, with select distances shown. The closest association was a single 7.6 A distance between TMPRSS2 Q438 and ACE2 K659. In the absence of SARS-CoV-2 RBD, TMPRSS2 was loosely associated at random locations, exemplified in Fig. 6C by a distance of 70.9 A separating TMPRSS2 active site residue H441 from each ACE2_K659 of the dimer-of-heterodimers.

**Fig. 6.**
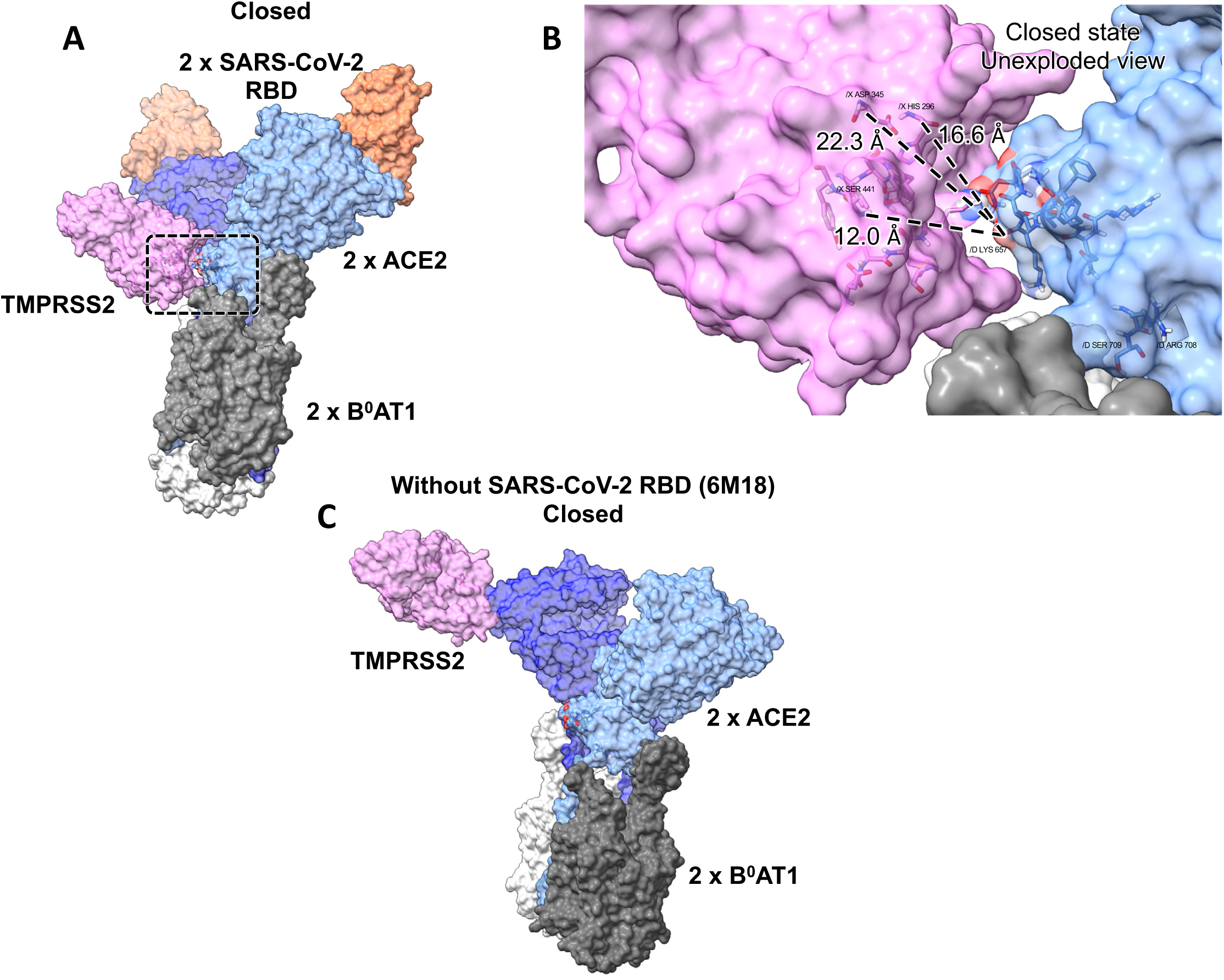
TMPRSS2 attempted failed contact with 2ACE2:2BOAT1 dimer-of-heterodimers complex. (A) Docking attempt results between TMPRSS2 and closed state (6M17) 2ACE2:2B^0^AT1 complexed dimer of heterodimers bound to two SARS-CoV-2 RBD. (ACE2, light and dark blue; SARS-CoV-2, light and dark gold; TMPRSS2, pink). (B) Closeup native view of non-exploded box in (A) showing lack of interface contact. ACE2 ectodomain putative cleavage zone residues were generally ∼15-30 A from TMPRSS2 active site pocket residues. The closest association was a single 7.6 A distance between TMPRSS2_Q438 and ACE2_K659. All attempted dimer-of-heterodimers docking models resulted in similar dysfunctional orientation random attachments. Details are in Table 1. (C) Results of TMPRSS2 ligand attempted docking with dimer-of-heterodimers lacking SARS-CoV-2 RBD (PBD ID: 6M18) under the same attraction/repulsion conditions as for (A). In this example of random loose attachment locations, the active site H441 residue of TMPRSS2 was loosely associated with K659 of one of the ACE2 subunits (light blue) at a distance of 70.9 A.

Fig. 7 and the data of Table 1 indicated that the open conformation of the 2ACE2:2B^0^AT1 dimer-of-heterodimers gave the same resulting failed docking with no interface contacts with TMPRSS2, as did the closed conformation of Fig. 6. In the open state, ACE2 ectodomain putative cleavage zone residues were generally ∼30 A from TMPRSS2 residues, as exemplified in Fig. 7B.

**Fig. 7.**
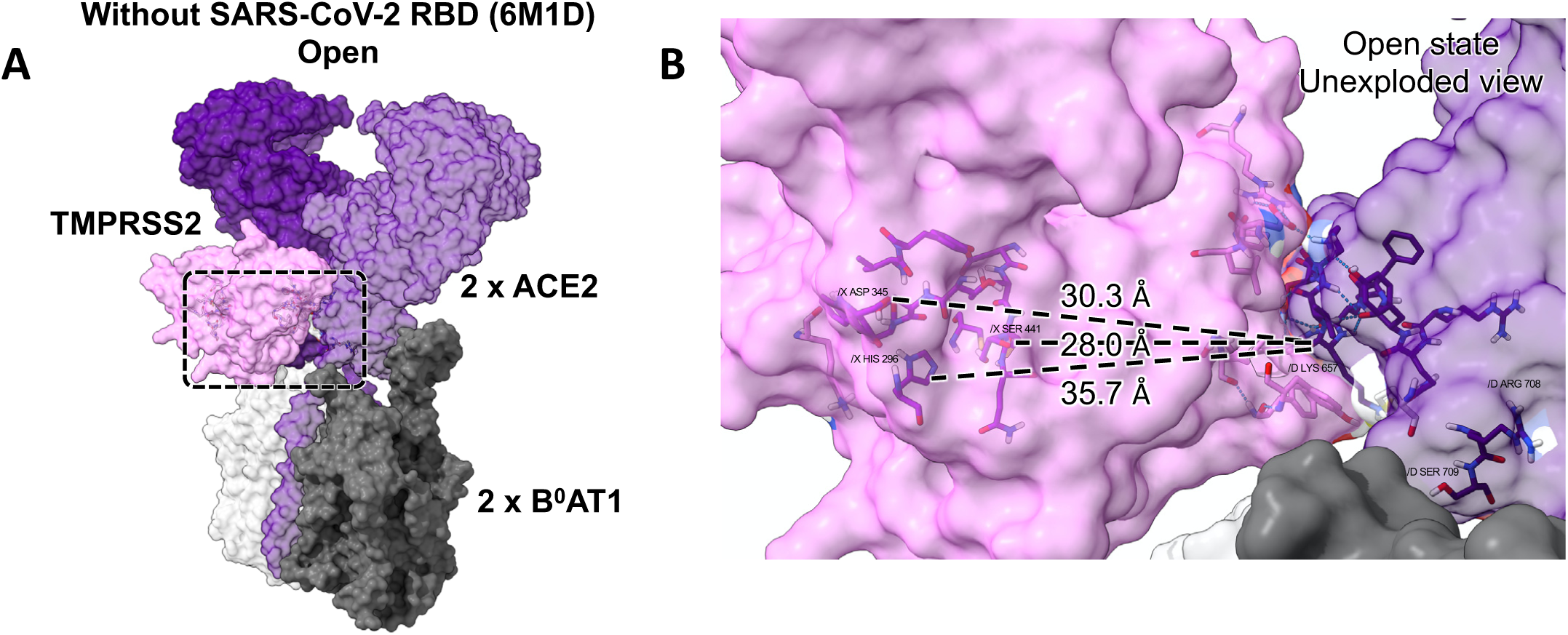
TMPR552 dysfunctional orientation random attachment to 2ACE2:2B0AT1 dimer-of-heterodimers open state complex. (A) Docking attempt results showing random attachment of TMPRSS2 to 6M1D open state 2ACE2:2B^0^AT1 complexed dimer of heterodimers. (ACE2, light and dark purple; TMPRSS2, pink). (B) Closeup native view of non-exploded box in (A) showing lack of interface contact. ACE2 ectodomain putative cleavage zone residues were generally ∼30 A from TMPRSS2 residues, with select distances shown. Details are in Table 1.

Fig. 8 shows successful docking of ADAM17 to stand-alone ACE2 monomer complexed with SARS-CoV-2 RBD, with details reported in Table 2. Residues of ADAM17 and ACE2 meeting criteria for interfacing “contact” are shown in Fig. 8 demonstrating participation of ACE2 neck region motif residues of Fig. 2B and 2C. An exploded separation of this contact zone emphasizing the active site catalytic pocket residues of ADAM17 with its zinc atom, is shown in the closeup in Fig. 8C with contact bond distances. Notably, ACE2_R710 engaged with a 1.77 A and 2.04 A double electrostatic attraction to ADAM17_H456, while equally notable was the lack of participation by ACE2_R708 which was 16.3 A from ADAM17_H415, and ACE2_S709 which was 17.3 A from ADAM17_H415.

**Table 2.**
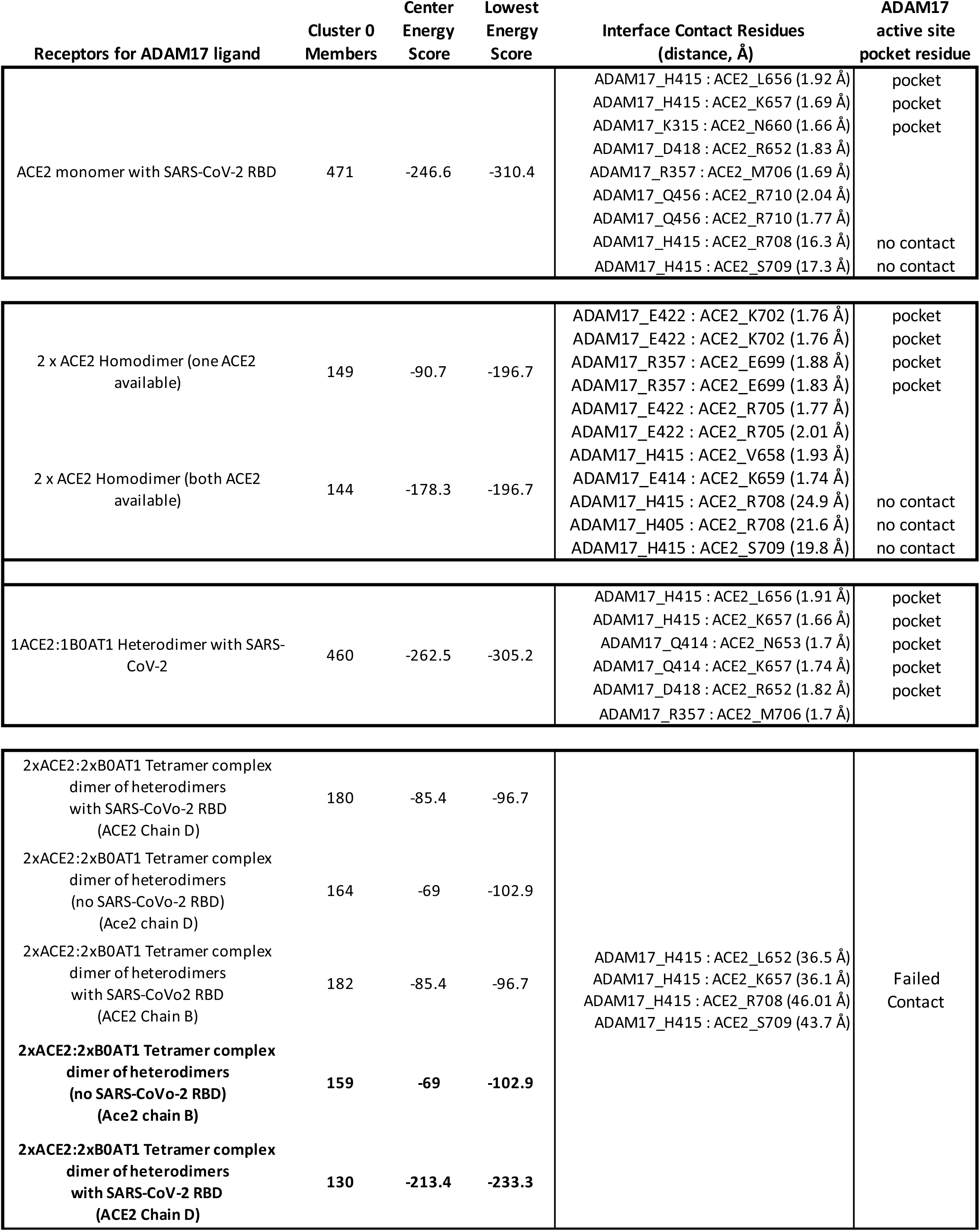
ADAM17 ligand molecular docking contact residues with various ACE2 structures.

**Fig. 8.**
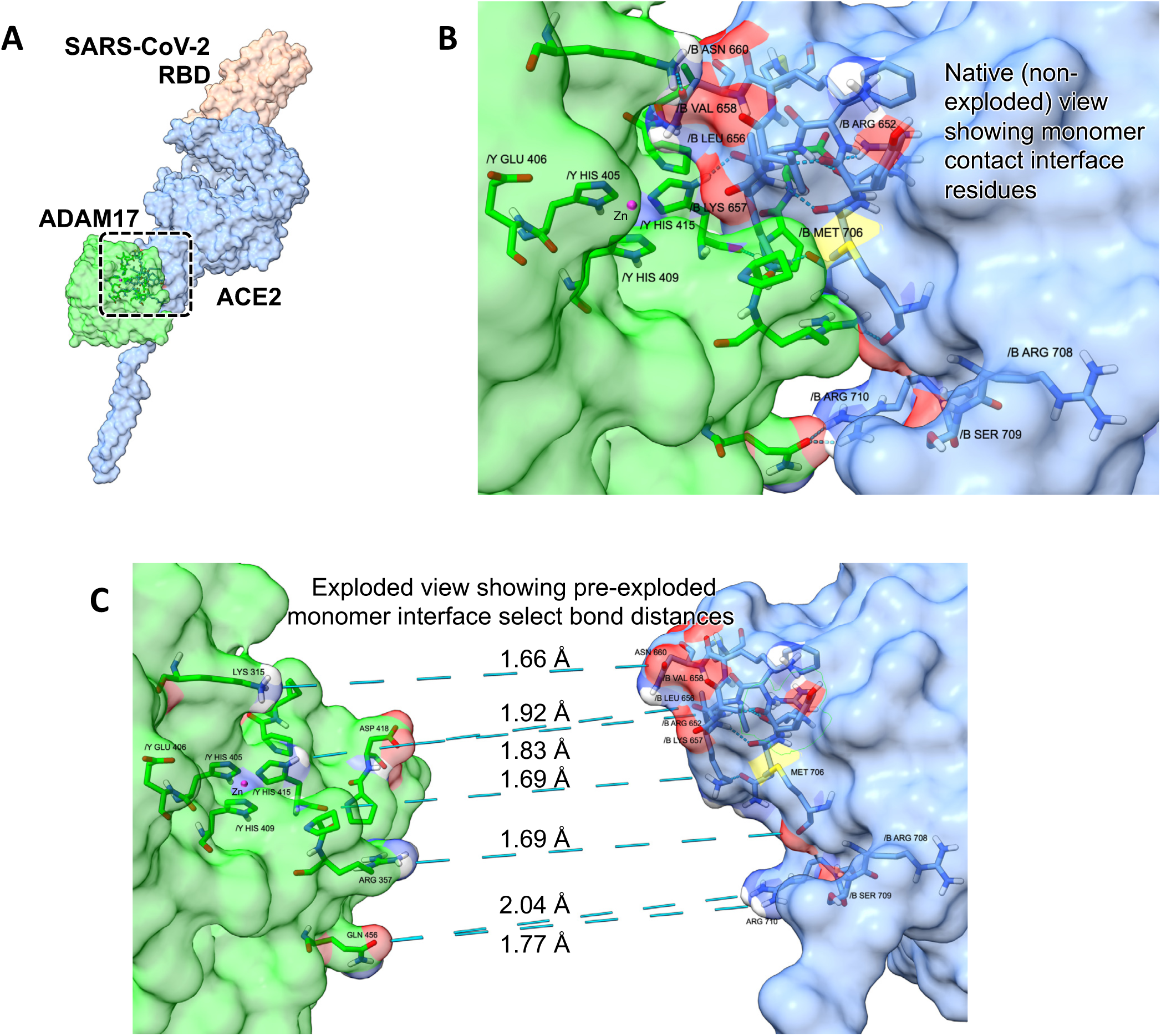
ADAM17 docked with ACE2 monomer complexed with SARS-CoV-2 RBD. (A) Docking model results showing ACE2 monomer (light blue), ADAM17 (green), and SARS-CoV-2 RBD (gold). (B) Closeup of box in (A) showing successful docking interface contact residues of ADAM17 catalytic active site pocket closely engaged with ACE2 ectodomain putative cleavage zone residues. (C) Exploded view of (B) native view showing pre-exploded interface contact select bond distances, with zinc atom (magenta)) in active site pocket. Select interface contact distances are shown, as measured using pre-exploded distances. Details in Table 2.

ADAM17 successfully docked with ACE2:ACE2 homodimer whether complexed with SARS-CoV-2 RBD (Fig. 9) or not including RBD (not shown). The resulting model interface residue contacts and distances are summarized in Table 2, which were the same whether docking conditions were restricted to availability of one or simultaneously both ACE2 subunit attractor residues. Fig. 9B shows an exploded view of the interface contacts between ADAM17 active site residues (with zinc atom) and ACE2 residues. Select distances are shown, with all bonds and distance interactions summarized in Table 2. Note lack of engagement of ACE2_R708 which was 24.9 A from ADAM17_H415, and ACE2_S709 which was 19.8 A from ADAM17_H415.

**Fig. 9.**
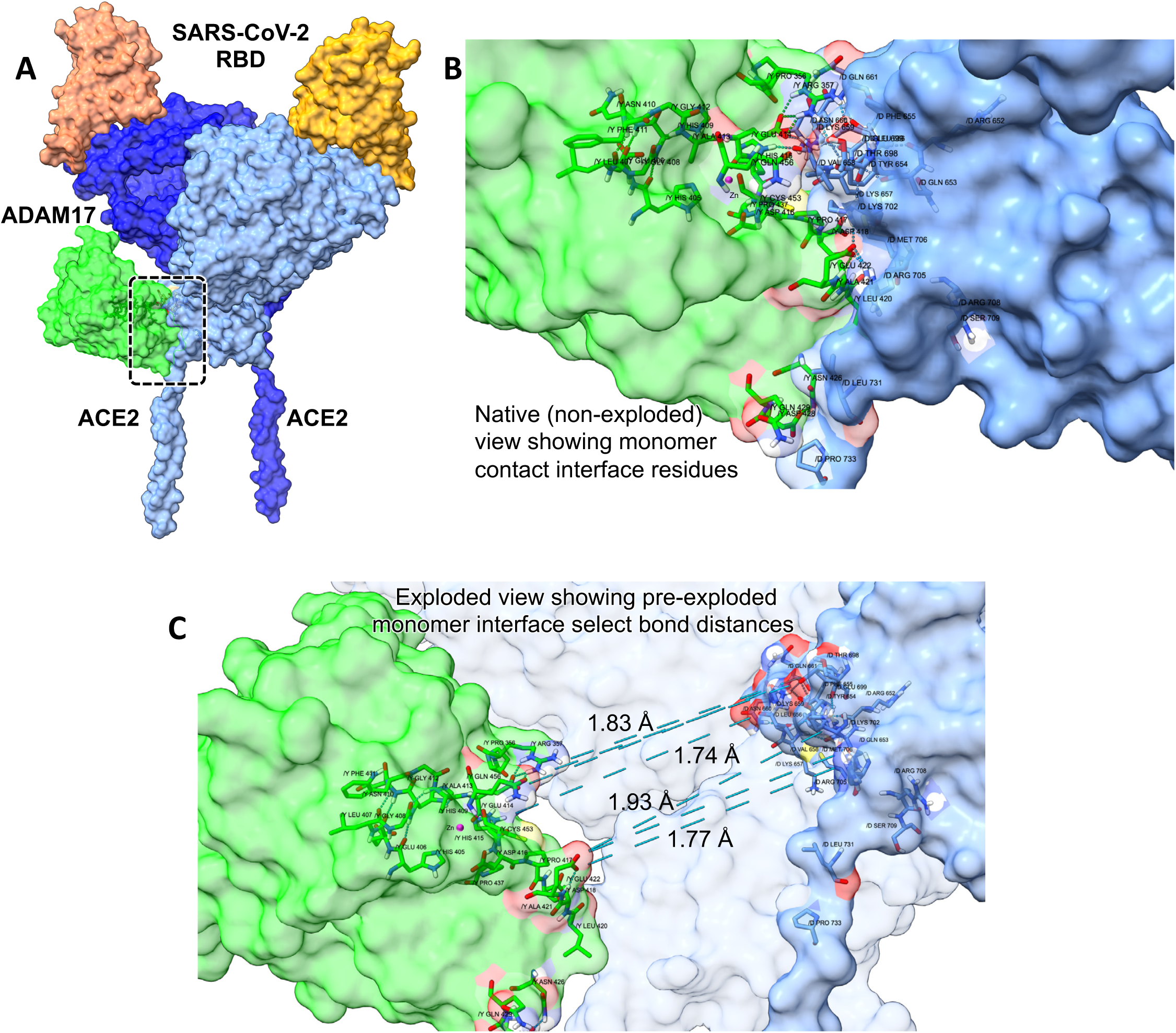
ADAM17 docked with ACE2:ACE2 homodimer complexed with SARS-CoV-2 RBD. (A) Docking model results showing ACE2 homodimer (light and dark blue), ADAM17 (green), and SARS-CoV-2 RBD (dark and light gold). (B) Exploded view of box in (A) showing docking interface contact residues of ADAM17 catalytic active site pocket (with zinc atom (magenta)) engaged with ACE2 ectodomain cleavage zone residues. Select interface contact distances are shown, as measured using pre-exploded distances. The same model results were obtained whether docking conditions were restricted to availability of one or simultaneously both ACE2 subunit attractor residues. Interaction details in Table 2.

The active site pocket of ADAM17 docked successfully with a heterodimer comprised of 1ACE2:1B^0^AT1 complexed with SARS-CoV-2 RBD, as shown in Fig. 10. An exploded view of the docking interface contact residues is shown in Fig. 10B along with select distances, with Table 2 reporting the bonds and distances.

**Fig. 10.**
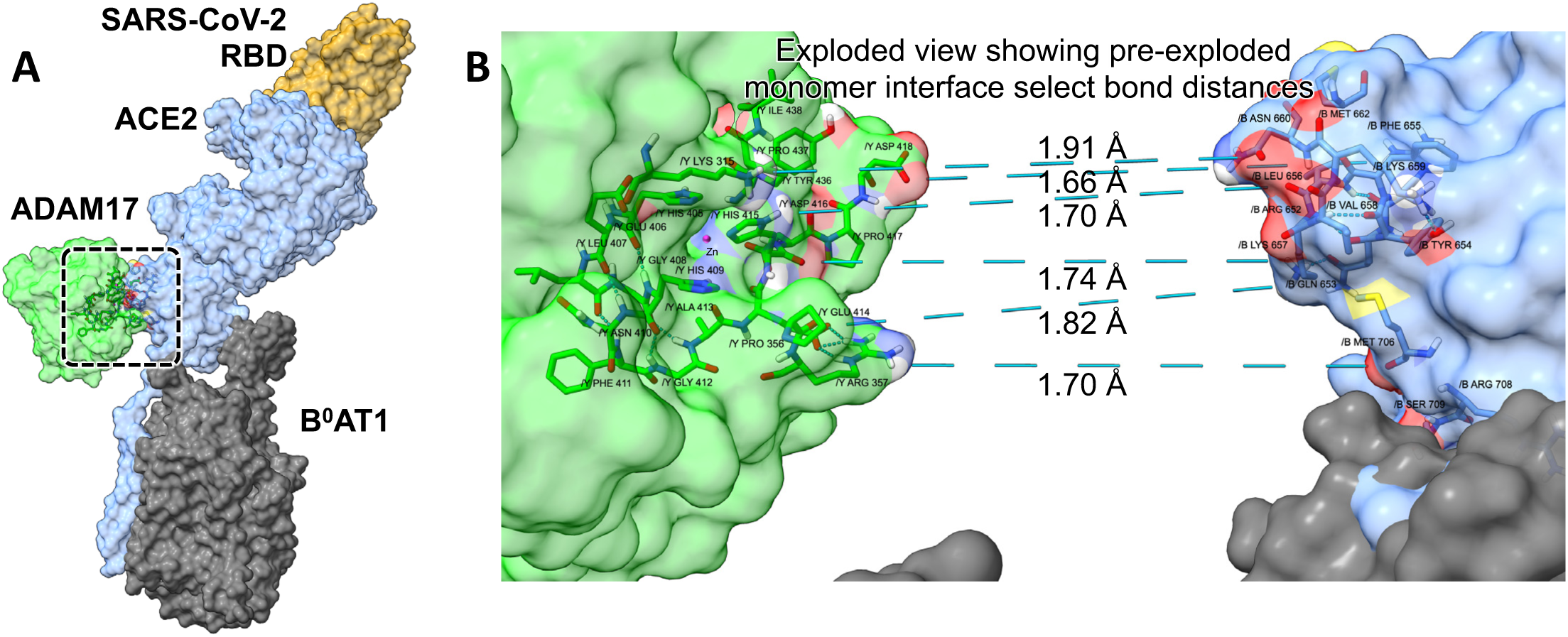
ADAM17 docked with 1ACE2:1B0AT1 heterodimer complexed with SARS-CoV-2 RBD. (A) Docking model results showing 1xACE2:1xB^0^AT1 heterodimer (ACE2, light blue; B^0^AT1, dark gray), ADAM17 (green), and SARS-CoV-2 RBD (gold). (B) Exploded view of box in (A) showing docking interface contact residues of ADAM17 catalytic active site pocket (with zinc atom (magenta)) closely engaged with ACE2 ectodomain cleavage zone residues. Select interface contact distances are shown, as measured using pre-exploded distances. The same model results were obtained whether docking conditions were restricted to availability of one or simultaneously both ACE2 subunit attractor residues. Interaction details are in Table 2.

Fig. 11 and the data of Table 2 indicated that ADAM17 ligand failed to dock with the closed state 2ACE2:2B^0^AT1 dimer-of-heterodimers neck region, whether or not the ACE2 was complexed to SARS-CoV-2 RBD. In repeated attempts employing various permutations of chain availability with or without RBD (Fig. 11A, Table 2), ADAM17 randomly stuck to the dimer-of-heterodimers in a dysfunctional orientation with its zinc atom and catalytic pocket structure residues outwardly facing 180^0^ away from ACE2 (Fig. 11B). In each case, active site ADAM17_H415 was generally ∼40 A from the dimer-of-heterodimers ACE2 neck region of Fig. 2.

**Fig. 11.**
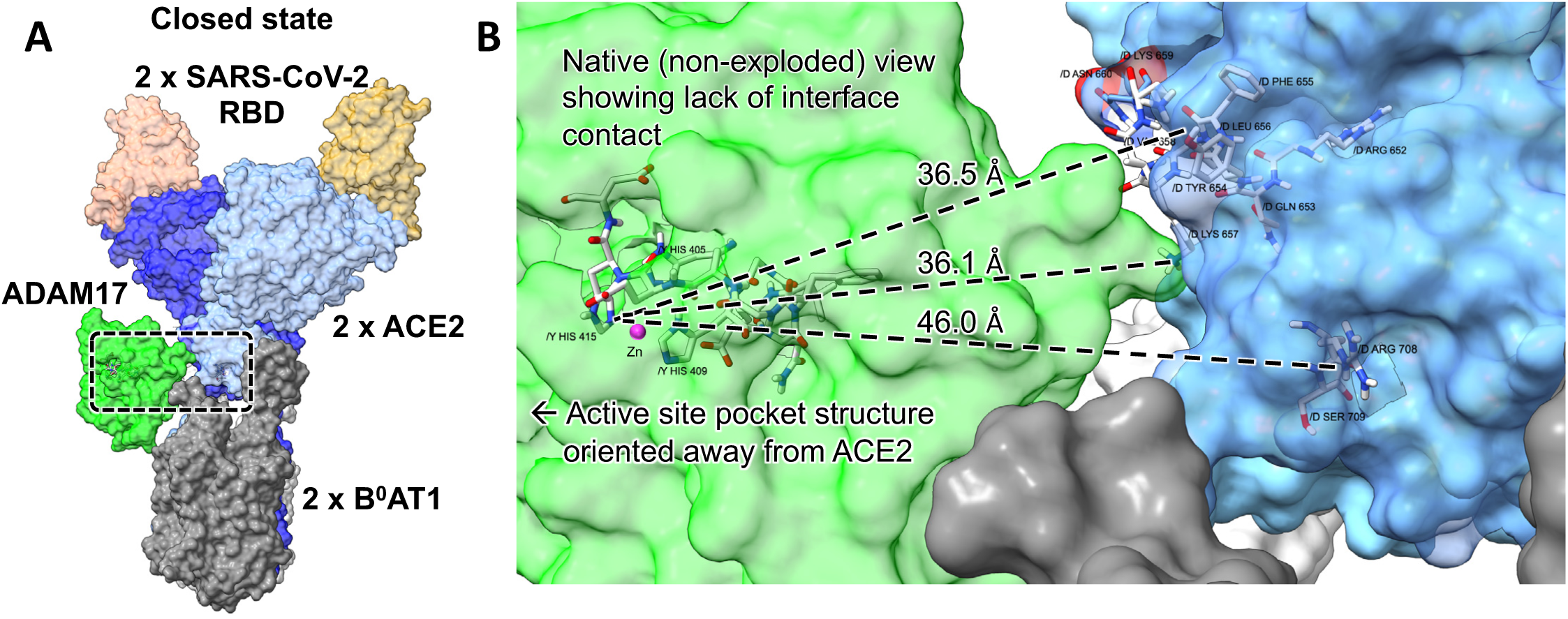
ADAM17 dysfunctional orientation random docking with 2ACE2:2B0AT1 dimer-of-heterodimers:SARS-CoV-2 RBD closed state complex. (A) Docking attempt results showing random attachment of ADAM17 to 6M18 closed state 2ACE2:2B^0^AT1 complexed dimer of heterodimers bound to two SARS-CoV-2 RBDs (ACE2, light and dark blue; SARS-CoV-2, light and dark gold; ADAM17, green with coordinated zinc atom, magenta). (B) Closeup non-exploded native view of box in (A) showing lack of interface contact. ACE2 ectodomain putative cleavage zone residues were generally ∼40 A from ADAM17_H415, with select distances shown. Note dysfunctional orientation of ADAM17 catalytic pocket structure with its active site residues outwardly facing 180^0^ away from ACE2. All attempted dimer-of-heterodimers docking models employing either SM17 (with RBD) or 6M18 (same as 6M17 but without RBD) resulted in similar dysfunctional orientation random attachments. The same results were obtained yielding models with similar cluster numbers and energies failing to dock with interface contact residues under all of the following conditions (not shown), whether: (i) SARS-CoC-2 RBD was present and attached (6M17 as in (A)) or not present nor attached (6M18); (ii) either of the ACE2 chains were individually targeted or (iii) both ACE2 chains attracted simultaneously. Details in Table 2.

Fig. 12 and the data of Table 2 indicated that the open conformation of the 2ACE2:2B^0^AT1 dimer-of-heterodimers gave the same resulting failed docking with no interface contacts with ADAM17, as did the closed conformation of Fig. 11. In the open state, ACE2 ectodomain putative cleavage zone residues were generally ∼40 A from active site ADAM17_H415, as exemplified in Fig. 12B, resulting in dysfunctional orientation of ADAM17 catalytic pocket structure with its zinc atom and residues of the active site pocket outwardly facing 180^0^ away from ACE2.

**Fig. 12.**
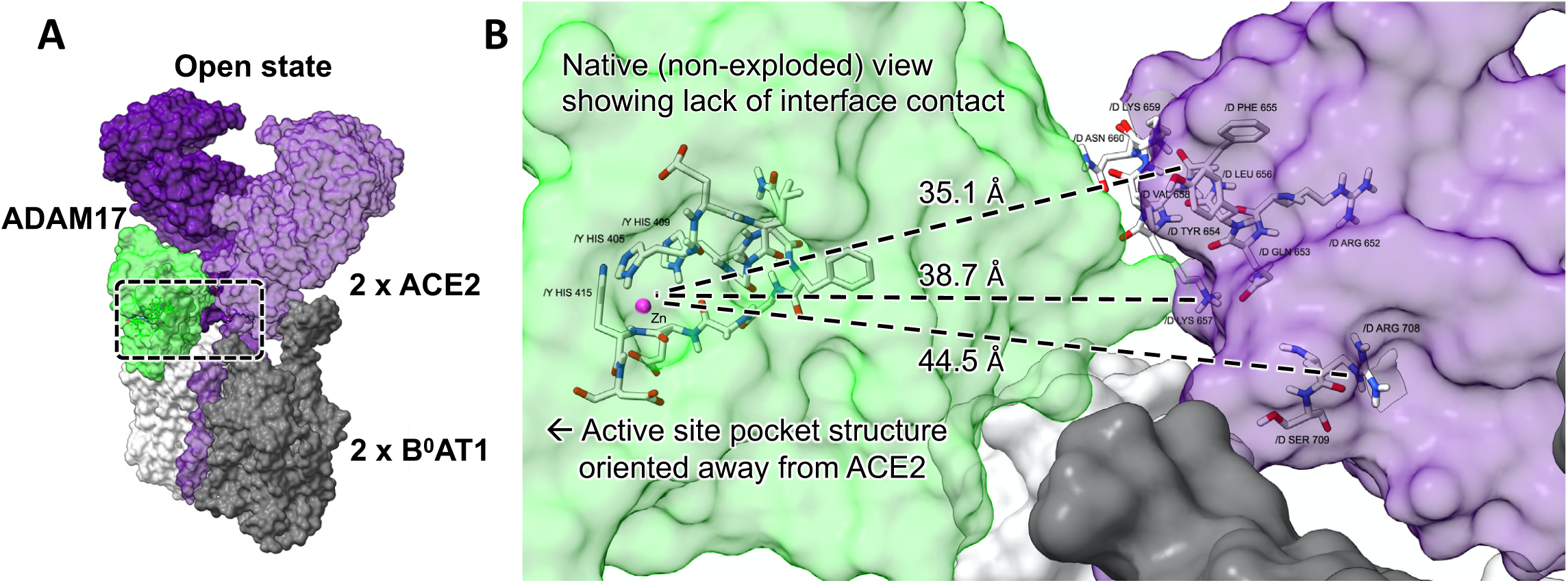
ADAM17 dysfunctional orientation random docking with 2ACE2:2B0AT1 dimer-of-heterodimers open state complex. (A) Docking attempt results showing random attachment of ADAM17 to 6M1D open state 2ACE2:2B^0^AT1 complexed dimer of heterodimers. (ACE2, light and dark purple; ADAM17, green with coordinated zinc atom, magenta). (B) Closeup of box in (A) showing native view (non-exploded) and lack of any interface contacts. ACE2 ectodomain putative cleavage zone residues were generally ∼40 A from ADAM17_H415, with select distances shown. Note dysfunctional orientation of ADAM17 catalytic pocket structure with its active site residues outward facing 180^0^ away from ACE2. All attempted dimer-of-heterodimers docking models resulted in similar dysfunctional orientation random attachments (not shown). Details are in Table 2.

## DISCUSSION

The principal finding is that the Results collectively implicate B^0^AT1 as a maJor putative player in the landscape of COVID-19 pathophysiology, especially in intestinal mucosa enterocytes, aging cardiomyocytes, and renal proximal tubule epithelial cells. These particular cell types share the unique common denominator kindred of expressing the B^0^AT1 neutral amino acid transporter in their plasma membranes [2, 5, 13, 14, 16-32]. B^0^AT1 was previously originally discovered and functionally characterized by Stevens et al. [21, 23-29] as the sodium-coupled neutral amino acid transport system in intestinal epithelial apical brush border membranes (alternately called NBB, B, B^0^) [21], with Broer, Fairweather, Verrey and colleagues [16, 18-20, 30-32] subsequently cloning its SLC6A19 gene and providing experimental evidence of trafficking/chaperoning of B^0^AT1 by ACE2. Yan and coworkers in Zhou’s group [17] employed cryo-electron microscopy to establish that the 2ACE2:2B^0^AT1 dimer of heterodimers is the thermodynamically favored atomic arrangement in either “closed” or “open” conformations.

Although pulmonary manifestations are the hallmark of COVID-19, SARS-CoV-2 exhibits experimentally and clinically reported gastrointestinal, cardiovascular and renal tropisms engaging local and systemic inflammasome pathophysiology that can appear before fever onset and extend after respiratory distress subsides [1-9, 13, 14, 40, 42].

The Results are further pertinent in comparing ACE2 monomer to the 2ACE2:2B^0^AT1 dimer-of-heterodimers. These are the most physiologically relevant states[13, 14], such that interpretations of biochemical experimental evidence primarily implicate existence of monomer ACE2 over homodimer ACE2:ACE2 in cell membranes not expressing B^0^AT1, and that the 2ACE2:2B^0^AT1 dimer-of-heterodimers is the most thermodynamically favored stabilized arrangement[17] in B^0^AT1-expressing intestinal membrane as shown by oocyte co-expression, immunoblots and native electrophoresis gels [2, 5, 13, 14, 16-32]. Nevertheless, in the interest of covering all possible permutations of docking combinations of ACE2 with or without B^0^AT1, the various dockings were attempted as reported in Results inasmuch as putative ACE2:ACE2 homodimer associations may occur under some circumstances, and the trafficking of ACE2 with B^0^AT1 could result in 1ACE2:1B^0^AT1 heterodimers under some circumstances. Furthermore, it is plausible that structural interactions of ACE2 with B^0^AT1 form a functional unit of peptide proteolysis and absorption of neutral amino acids in intestinal or proximal tubule epithelial luminal membranes.

In each of these gut, cardiac and renal cell types, ACE2 is known to indispensably counterbalances the pernicious arm of local renin-angiotensin system (RAS) as a native function [13, 14, 18, 31, 32, 40, 42, 44, 45, 47, 53, 67-76], and is the hiJacked receptor of SARS-CoV-2. In each of these cell types along with pneumocytes, TMPRSS2 and ADAM17 are prominent proteases expressed with ambiguous benefits but many pathological downsides. ACE2, TMPRSS2 and ADAM17 engage in an untoward triangulation known to drive various disease states involving spiraling positive feedback unchecked tissue behaviors and inflammation, notably in COVID-19 [1-3, 12, 44, 77, 78]. Critical structural and physiological interactive relationships exist among the players ADAM17, dimer-of-heterodimers 2ACE2:2B^0^AT1, and SARS-CoV-2 receptor binding domain (RBD) of ACE2, but this interrelationship is largely unexplored and poorly understood.

As demonstrated by the dockings of various ACE2 arrangements each involving residues of the neck region (Fig. 2), the Results Figs. 3, 4, 5, and Table 1 show that the TMPRSS2 active site pocket catalytic triad H296, D345 and S441 and the substrate binding domain residues D435, S460 and G462 concertedly formed contact bonds of the appropriate lengths aligned in the correct three dimensional orientation that has been experimentally determined to fit known TMPRSS2 substrates [49-51, 64]. In concert with our validation of the steric arrangement of the TMPRSS2 active site pocket residues (Fig. 1), the three dimensional structure employed in the present proJect has been successfully used previously as a TMPRSS2 surrogate for drug discovery by way of its strong docking with steric accuracy to a variety of known classic drug agonists and antagonists of TMPRSS2 [49-51, 64]. SARS-CoV-2 infectivity in human cells can be inhibited by Camostat, Nafamostat and several other experimental small molecules[49-52]. These strongly interact with TMPRSS2’s active site pocket involving the catalytic triad of H296, S441 and D345 and substrate recognition residues D435, S460 and G462 [49, 50, 64, 79, 80]. Reported molecular dynamics docking binding energies for this structure of TMPRSS2 include −7.20 kcal/mol [51] or −7.94 kcal/mol [50] for Camostat mesylate, and −7.21 Kcal/mol [51] or −8.20 kcal/mol [50] for Nafamostat, and the highest molecular docking score for experimental anti-SARS-CoV-2 compound NPC306344 [64]. The data in Table 1 and Figs. 3-5 show that TMPRSS2 docking model clustering, energy minimizations, and ligand:target interface residue contacts were virtually the same for ACE2 whether as a monomer or as a 1ACE2:1B^0^AT1 heterodimer. Docking of TMPRSS2 with the ACE2:ACE2 homodimer conformation was also successful, although clustering, energy scores and residue participations (Table 1) were not as robust as monomer or heterodimer. In stark contrast, the data of Table 1 and Figs. 5-7 show that 2ACE2:2B^0^AT1 dimer-of-heterodimers failed to dock with TMPRSS2 within precepts of physiological reality under any simulation conditions. Here, the failure of TMPRSS2 docking and lack of active site pocket participation held for any simulation condition, whether the dimer-of-heterodimers was in the open or closed conformation, with or without SARS-CoV-2 RBD attached.

ADAM17 is a member of the metzincins which all possess the catalytic HEXXHXXGXXHD motif with a zinc atom, and in particular for the present study the catalytic pocket of ADAM17 PDB ID:3LGP chain A was HELGHNFGAEHD [66]. Results Fig. 8 and the data of Table 2 indicate that monomeric ACE2 strongly docked with these ADAM17 active site pocket residues with steric accuracy at H405, E406, L407, G408, H409, N410, F411, G412, A413, E414, H415, and D416 engaging the zinc coordinated catalytic triad of H405, H409, H415 in conJunction with glutamate at E406 serving an acid/base catalytic function [66]. The Results Fig. 9, Fig. 10, and Table 2 data showed that the ADAM17 active site pocket also engaged appropriately and strongly with the ACE2:ACE2 homodimer or 1ACE2:1B^0^AT1 heterodimer models. The Results indicated that ADAM17 docking clustering and energy minimization scores and ligand:target interface residue contacts were nearly the same for ACE2 monomer and 1ACE2:1B^0^AT1 heterodimer, both of which were quantitatively more robust than the successful docking with ACE2:ACE2 homodimer. The monomer, homodimer, and heterodimer results shown for ACE2 complexed with SARS-CoV-2 were also each the same if SARS-CoV-2 RBD was not attached to ACE2 (data not shown). Contrasting these successful dockings, the data of Table 2 and Figs. 8-12 showed that ADAM17 failed to dock with 2ACE2:2B^0^AT1 dimer-of-heterodimers with or without SARS-CoV-2 RBD, and whether or not in the “open” or “closed” conformational state. Indeed, modeling results showed ADAM17 randomly bumping into the 2ACE2:2B^0^AT1 dimer-of-heterodimers with the active site pocket structure dysfunctionally orientated outward facing 180^0^ away from ACE2, with the ADAM17 catalytic triad residues ∼40 A distant from ACE2 (Figs 8-12, Table 2).

Based on biochemical experimental data derived using transgenic expression systems, Heurich et al. [47] implicated ACE2 residues 652-659 as essential for binding recognition by TMPRSS2 or ADAM17, but not necessarily the cleavage site. The Heurich experiments [47] further pointed to the ACE2 ectodomain shedding putative cleavage site as likely within its neck region nebulously somewhere in the span of residues 697-716. Also employing biochemical experiments, Jia et al [81] contended that the putative cleavage might occur in the span of ACE2 residues 716-741, while Lai [53] implicated cleavage between residues R708/S709 with 710 playing a role in presumed binding recognition.

The present docking modeling Results yielded mixed agreements with these prior [47, 53, 81] experimental suggestions of putatively important residues. The docking data of Tables 1 and 2, and Figs. 1-12 indicated that residues spanning R652-D713 in ACE2 of all its forms (except in 2ACE2:2B^0^AT1 dimer-of-heterodimers) made important interface contacts with each TMPRSS2 and ADAM17, with notable prominence of ACE2 N660 in either monomeric, homodimeric or heterodimeric formats participating in multiple short distance bindings with key residues of TMPRSS2 (H296, S441, G462, S460), and to a lesser extent to ADAM17 residue K315. The docking Results further showed that the active site pockets of TMPRSS2 and ADAM17 each formed strong multi-bond interfaces with ACE2 K657, and that monomer and heterodimer ACE2 M662 participated in bonds with pocket residue G462 of TMPRSS2. Furthermore the docking results indicated that: i) ACE2 R710 provided electrostatic attractions to ADAM17 Q456 (1.77 and 2.04 A double bonding) in the ACE2 monomer, or to TMPRSS2 E299 with the ACE2 heterodimer (1.82 A); ii) monomeric or heterodimer ACE2 S709 formed a double bond attraction (ranging 1.71-1.88 A) with non-catalytic K300 of TMPRSS2; but iii) S709 in none of the ACE2 arrangements formed an interface contact with any ADAM17 docking (distances ranging 17.30-43.70 A); and i) ACE2 S708 in any arrangement never met interface contact criteria for, nor even proximity to, any catalytic pocket residues of either TMPRSS2 nor ADAM17 (distances ranging 16.30 − 46.01 A).

In conclusion, the present molecular docking observations implicate the neutral amino acid transporter B^0^AT1 (SLC6A19) [2, 5, 13, 14, 16-32] as a maJor player in governing TMPRSS2 and ADAM17 steering of ACE2’s multifaceted roles in COVID-19, whether in the realm of modulating ACE2’s role in RAS and inflammation, its roles in interactions with SARS-CoV-2 binding and infectivity, or in considerations of cross-interferences

